# An integrative resource for investigating genetic model of drug response in cancer

**DOI:** 10.1101/555789

**Authors:** Yi Guo, Xiang Chen

## Abstract

**Background:** The most important goal of pharmacogenetics research is to obtain genetic models closely related to drug efficacy *in vivo* and *in vitro*, which can be used to measure individual responses and then to assess the sensitivity and toxicity of anticancer drugs. However, these models are scattered in different pharmacogenetic studies and are difficult to be effectively used and analyzed.

**Results:** To this end, we have developed an integrated resource named iGMDR that provides the genetic models of both clinical and preclinical anticancer drugs. iGMDR provides a unique resource for mining the links between anticancer drugs and individual genomes, which will attempt to discover cancer knowledge and unlock the potential value of this data.

**Conclusions:** We developed iGMDR as a web service whose code was released at GitHub repository of ModelLAB (https://github.com/ModelLAB-ZJU/iGMDR), it also provided online access and application interface (https://igmdr.modellab.cn). MySQL dump for the integrated resources was stored at zenodo.org (DOI: 10.5281/ zencomod.1432561).

## BACKGROUND

Pharmacogenetics (a portmanteau of pharmacology and genetics) is the study of the role of genetics in drug response, and the main goal is to find more efficient disease treatment strategies based on individual genetic characteristics [1, 2]. It is one of the main bottlenecks in the implementation of personalized medicine due to the complexity of research [3, 4]. Traditionally, diseases have been treated largely based on an epigenetic understanding and a knowledge of population genetics of the disease, and results often fail in specific individual cases [5]. With the rapid development of genetics-related sequencing technology and the accumulation of knowledge due to various drug development, the relationship between the curative efficacy and the individual genetic characteristics has become increasingly clear, leading to more accurate prediction of the effects of treatment strategies based on the genetic characteristics of patients, and thus we have obtained genetic models of some drugs with widespread clinical application [3]. Cancer therapy is one of the challenging goals of precision medicine and personalized medicine, and its occurrence is closely related to abnormal regulation of cellular function level (such as genome, transcriptome and proteome events) [6]. As a result, pharmacogenetics studies in cancer have been widely implemented and have produced many effective genetic models that have been applied in clinical practice (NCCN guidelines, FDA drug labels) [7, 8]. Nevertheless, the pace of research in cancer pharmacogenetics has lagged far behind the need for cancer precision medicine, and many cases of access to treatment remain based on traditional “one treatment fits all” strategies [9]. More systematic studies and more effective data analysis are needed if more individual-specific genetic models are to be obtained to improve the effectiveness of cancer treatment. Experimental designs from *in vitro* to *in vivo* are an indispensable path for the application of majority knowledge to clinical practice. Currently, there are several *in vitro* pharmacogenetics studies based on cancer cell lines, such as CCLE [10], GDSC [11], CTRP [12], CGP [13], and MCLP [14], and these studies have yielded many genetic models for specific drugs and specific types of cancer. In addition, *in vivo* cancer studies of some model organisms have also yielded considerable preclinical genetic models. While there are still many gaps and unpredictable challenges to translate these genetic models into clinical practice, the value of these data should not be underestimated. To this end, we constructed iGMDR to collect genetic models of anticancer drugs in different technical systems and different research stages. Our research goal is to integrate big data content and create new value based on these data.

## DATA CURATION AND ACCESSION

Recently, iGMDR integrates datasets including ClinicalTrials, FDA, NCCN, ASCO (https://www.asco.org), CCLE, GDSC, CTRP, MCLP, CGP, and literature on pharmacogenetic studies. As described above, we collected genetic models of anticancer drugs from four sources: 1) clinical guidelines, 2) approved drug labels, 3) scientific literature, and 4) experimental statistics (see Figure 1, for detailed reference datasets on the online website). According to the identification stage, it can be divided into clinical and preclinical models; According to the research object, it can be divided into *in vivo* model and *in vitro* model; Genome, transcriptome, proteome model and logical model (logical combination of events) were obtained according to the classification of the technical system. According to the model related event types mainly include SNV (simple nucleotide variation), CNV (copy number variation), EXP (expression), SV (structural variation), SPV (splice variant) and LN (cell lineage) [see Table S1 for more event types]. We standardized the data from these sources, including tissue type, cancer type, drug, gene and model presentation (see Table S2 for detailed process description). Eventually, we obtained about 154,100 models of 144 cancers of 30 tissue types, involving 1,040 different types of anticancer drugs (hormones, small molecules, vaccines, antibody drugs, etc.). In order to build an efficient research resource of pharmacogenetics, we integrated various information about drugs including chemical composition, structure, target, signal pathway and classification, and information about genes includes functional description and associated functional signaling pathways (see Table S3 for a list of detailed information sources).

**Figure 1.**
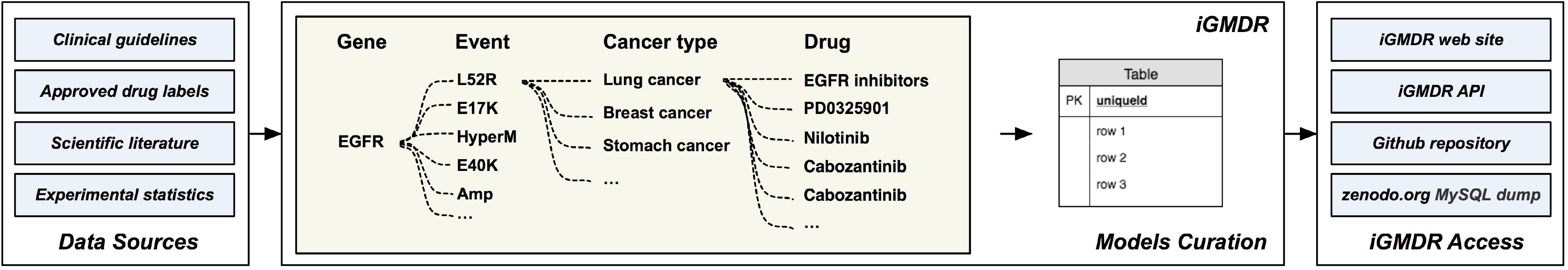
The schematic overview of iGMDR. The first column describes the data sources of the genetic model of anticancer drugs. The second column describes the data integration process. The third column is how the data is accessed and retrieved.

## WEBSERVER CONSTRUCTION

Based on the above integration process, we constructed an anticancer pharmacogenetics model resource, iGMDR, whose code is open source and available on GitHub repository (https://github.com/ModelLAB-ZJU/iGMDR) based on the GPLv3 license. The iGMDR online site was built on an Apache HTTP server, all model data was stored in a MySQL database, and PHP was used for backstage and front-end interactions (see Figure 1). Users can fully browse and download the genetic models of anticancer drugs through different data types (cancer, tissue, drug, and gene). Based on the intuitive understanding that pharmacogenetics researchers are usually interested in genetic models involving specific genes and anticancer drugs, we also designed the relevant search and profile interfaces, and the feedback information will be displayed visually. Specific drugs exhibitions will include general information (composition, targets, signaling pathways, classification, and external database ID), conformer structure (small molecule drugs), models of drug-related genes, the cancer tissue distribution of drug-related models, the signaling pathways and function enrichment of drug-related models, the drug-gene interaction network (the network was built through the number of drug-related models), and the drug-related model list. Specific gene profiling page describes the basic information of the gene (gene summary, categories, and external database ID), gene-related signaling pathways and functions, gene-associated anticancer drugs, the normal tissue distribution of gene expression in human, the distribution of gene-related drug targets and signaling pathway, and gene-drug interaction network (constructed by the number of gene-related models). These profiling and data presentations will facilitate the understanding of specific anticancer drug and gene-related genetic models.

## IMPLEMENTATION CASE

Lung cancer is one of the most popular cancers in the world [15]. Over 7.15% (COSMIC) of cancer patients have the defects (Point Mutations, Copy Number Variation, and Gene Expression) of STK11 gene. STK11 is a tumor suppressor that functions in the regulation of cell polarity, TP53 activity, and cell-cycle arrest. In our integrated resources, we can find that the most influential cancer tissue related to STK11 is Lung. For Lung cancers harboring STK11 mutations, we can predict the intervention effect of the corresponding 113 drugs. Of course, the genome variation of cancer patients is complex and diverse, and the genetic indicator from a single model may not be able to predict the final efficacy of drugs, so our resources also provide other indicators of genetic variation to predict the intervention effect of these 113 drugs.

Mahoney et al. have reported that non-small cell lung cancer (NSCLC) cell lines with mutations of STK11 and KRAS genes are sensitive to MEK inhibitors showing a dose-dependent inhibition of growth rate, while when the two are mutated alone that will show opposite inhibition [16]. In addition, several groups of clinical trials (e.g. PF-4554878, VS-6063) recruiting cancer patients for NSCLC have been recommended to test the status of STK11 [17, 18]. As shown in Figure 2, we constructed a gene-related model network of STK11 in our resources, including all the genetic indicators of these 113 drugs (red dots). Importantly, these drug interventions for NSCLC can be found in this network. With the help of this global model network, treatment strategies can be tailored to a patient’s own genetic defects.

**Figure 2.**
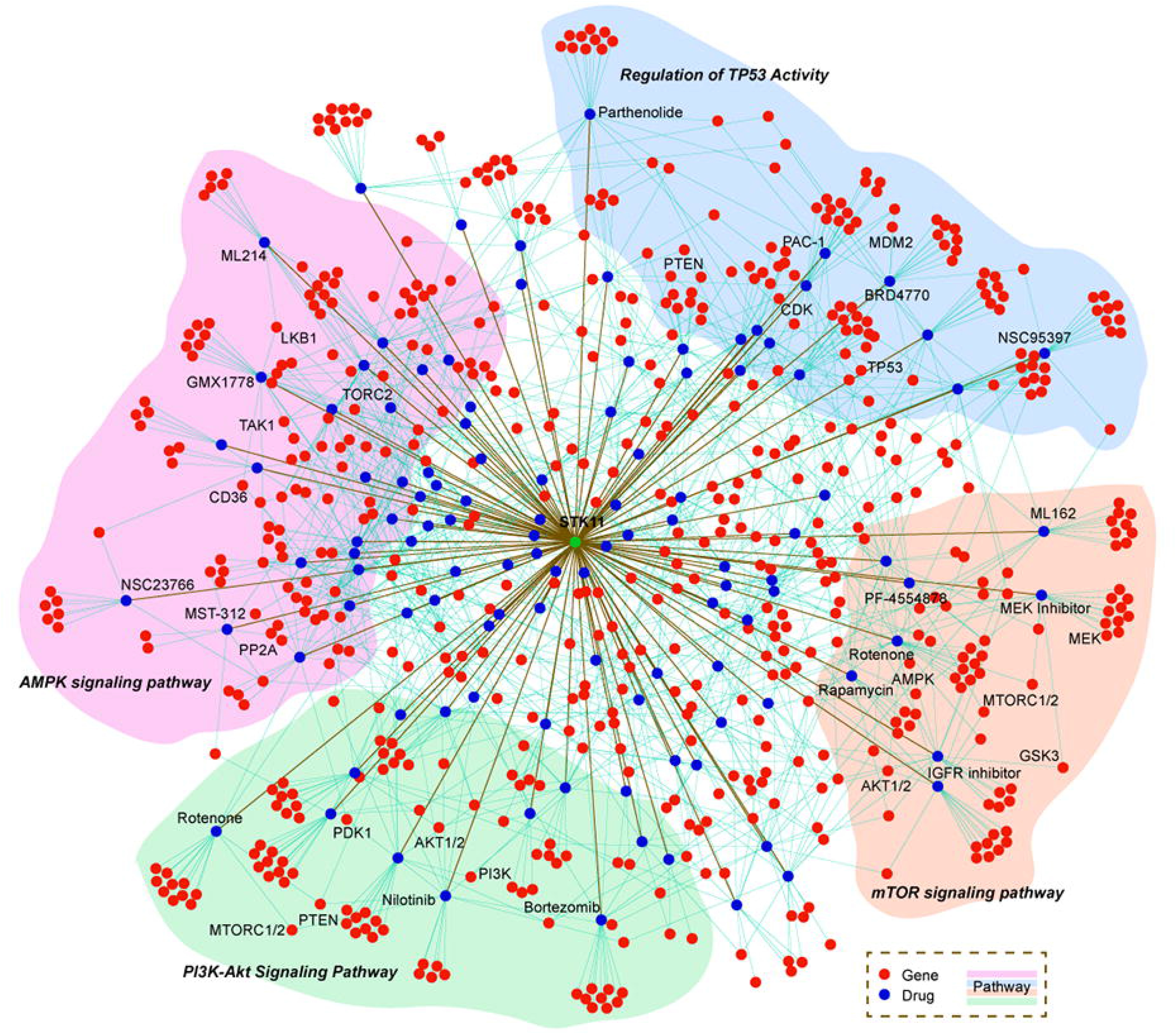
STK11 gene related drug model (drug-gene) network. Model related drugs and genes were connected by straight lines, while drugs directly related to STK11 genes were connected by significantly thick lines. The background color blocks represent the signaling pathways of the genes associated with these models.

The essence of tumorigenesis has been widely believed to be the result of aberrant cell signal transduction caused by genetic variation that further disrupt these signaling pathways as cancer progresses, and this process will become much more complicated than that of tumorigenesis [19]. Therefore, the traditional targeted therapy for a single target is often less effective for cancer treatment. Through the query of our resources, multiple targeted drugs are sensitive in terms of Lung cancer with STK11 gene defects, and the genes targeted by these therapeutic drugs participate in different signaling pathways (see Table S4). In addition, the predictive genes involved in these drug-related treatment models are also involved in these different signaling pathways (Figure 2). Based on this, we can design new treatment schemes to deal with the same or different pathways with abnormal signal transduction to improve the therapeutic effect of cancer. In other words, we can use this network to further infer new pharmaceutical strategies.

Interestingly, some kind of strategy at the signaling pathway level and important inferences for cancer treatment had been really reflected in the model networks that we built on this resource. STK11 is frequently mutated in NSCLC that acts as a tumor suppressor by activating AMPK (5’ AMP-activated protein kinase) signaling whereas loss of STK11 by point mutation or deletion suppresses AMPK, leading to increased mTOR signaling. Recently researchers have investigated the effects of STK11 mutation and mTOR inhibition on cell signaling pathways in NSCLC cell lines, suggesting a feedback mechanism that STK11 mutant cell lines increased insulin-like growth factor receptor (IGFR) activity, and inhibition of the IGFR activity was shown to downregulate the mTOR pathway[20]. The results supported the investigation of IGFR inhibitors in combination with drugs targeting the mTOR pathway, particularly for tumors bearing STK11 alterations [21].

By dephosphorylation of PIP3, PTEN negatively regulates PI3K/AKT leading to tumor suppression in NSCLC [22]. PTEN inactivation/loss therefore increases activity of the PI3K-AKT-mTOR pathway and further suggest that NSCLC patients might afford an opportunity for exploitation of anti-PI3K-AKT/mTOR-targeted therapies. Several studies have shown this subset of lung cancers treated with PI3K/AKT inhibitors combining with mTOR inhibitors (rapamycin) [16]. Until now, targeting of TP53 signaling has proved to be highly disappointing. However, recent studies indicate abundant crosstalk between TP53 and the AKT/mTORC1 signaling pathway can determine the choice of response to TP53. These results could have important pharmacological consequences for the potential development of PI3K/Akt inhibitors against TP53-deficient cancers [23, 24].

To sum up, the integrated model information will provide guidance for personalized treatment of cancer patients, and users can also infer new therapeutic strategies by connecting the model with the signal transduction network so as to improve the defects of single-targeted treatment (see Figure 3). These directions may help to identify patients who would most benefit from alternative single or dual pathway inhibition potentially leading to a revision in current molecular testing guidelines, and also demonstrate the advantages of integrating resources.

**Figure 3.**
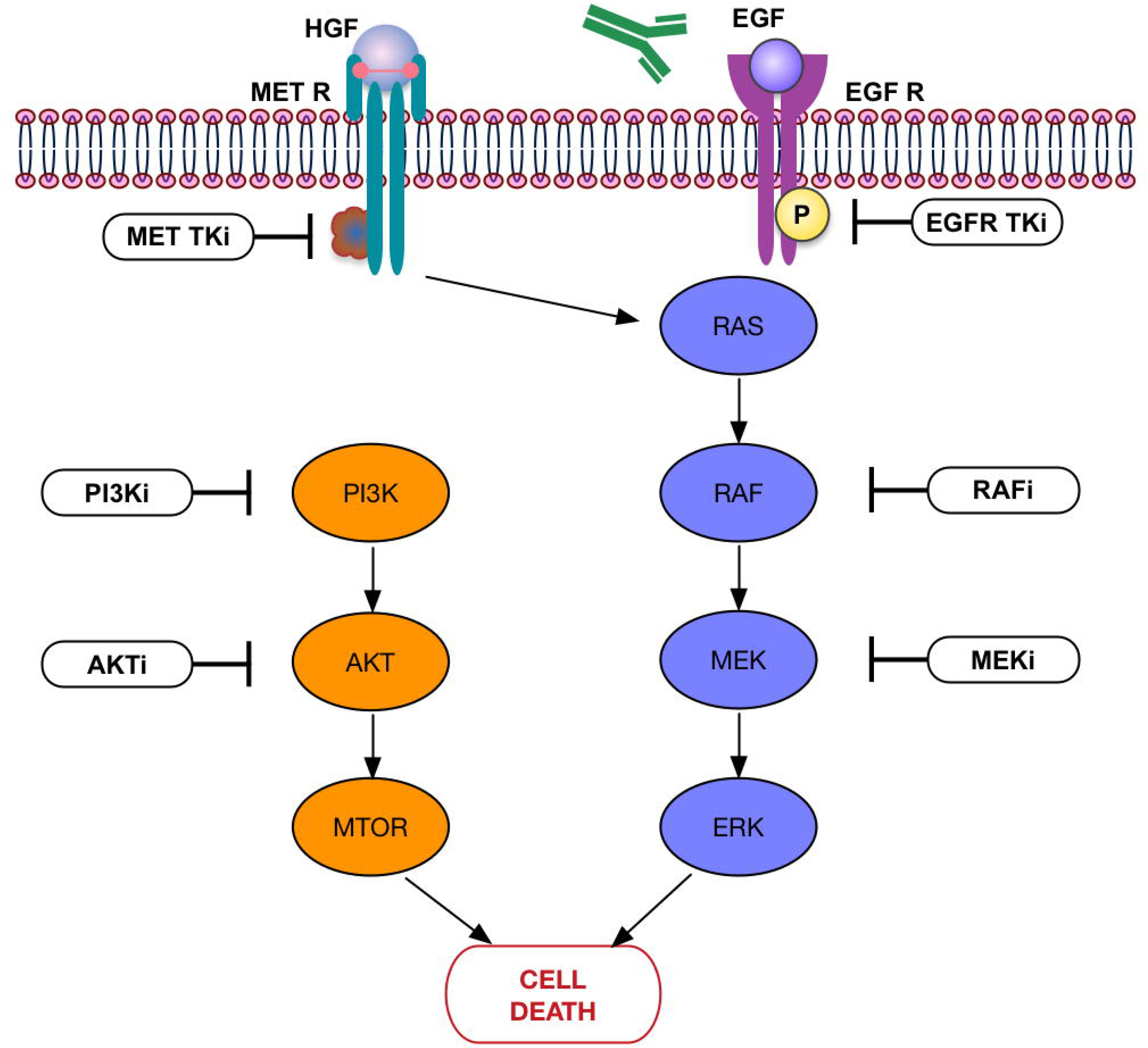
Mechanism diagram of cancer treatment strategy based on cellular signaling pathways. This diagram simulates the processes of drug intervention against abnormal signaling pathways in cancer cells.

## SUMMARY AND APPLICATION DIRECTIONS

Drug genetics model can effectively achieve precision anticancer treatment [25, 26]. To release the value of existing data, we have established a draft principle to collect the drug genetics model. iGMDR is the first integrated data resource to provide a predictive model for the therapeutic effect of anticancer drugs. Compared with the resources we have integrated, the current resources only provide a single genetics model of anticancer drug (“experimental statistics” for CCLE and GDSC or “clinical guidelines” for NCCN guidelines and FDA drug labels), and the data volume is also different by orders of magnitude, which greatly limits the application value of such data. The interactive design and data visualization of the iGMDR website provides both macro and micro insights for pharmacogenetics researchers in the cancer drug genetics model. For example, we can design a new sequencing panel for cancer drug efficacy prediction; We can link drug-related targets and signaling pathways to design combinatorial strategies for cancer therapy. In conclusion, iGMDR will continue to integrate relevant pharmacogenetic studies into cancer clinical and research therapy.

## Competing interests

The authors declare that they have no competing interests.

