## supplementary information for "An integrative resource for investigating genetic model of drug response in cancer"

**Table S1.** Event descriptors and descriptions in iGMDR models.

**Table S2.** Normalized organization of tissue, cancer types, anticancer drugs and models.

**Table S3.** Annotation of model-related drugs and genes with external resources.

**Table S4.** STK11-related genetic models involve drug targeting signaling pathways. Abnormalities in these signaling pathways are key players in the development of many cancer types. For different pathways, the table lists the number of drugs involved in the signaling pathway and the number of drug-related models.

**Table S1.**

| **Event Descriptor** | **Description** | **Example** |
| --- | --- | --- |
| AMP | amplification | amplification; CNV-H |
| BIA-IA | biallelic inactivation | biallelic inactivation |
| CNV | copy number variation | CNV |
| DEL | deletion | deletion; copy-neutral loss of heterozygosity; CNV-L |
| DEL-MUT | deleterious mutation | deleterious mutation |
| DEL-PLM | deletion polymorphism | deletion polymorphism |
| D-MUT | domain mutation | exon nucleur domain mutation; tyrosine kinase domain mutations |
| EXP | expression mutation | expression; serum levels |
| FL | function loss | any function loss |
| FS-MUT | frameshift mutation | frameshift mutation |
| FUS | fusion | fusion; translocation; rearrangement |
| GT | genotype | OncoGeno; HOMOZYGOSITY |
| HM | hypermethylation | promoter hypermethylation |
| IF-DEL | inframe deletion | inframe deletion |
| IF-INT | inframe insertion | inframe insertion |
| INDEL | insertion/deletion | insertion/deletion |
| INT | insertion | insertion; InsT |
| ISOF | isoform | isoform; alternative transcript; ISOFORM EXPRESSION |
| ITD | internal tandem duplication | internal tandem duplication; TANDEM REPEAT |
| LN | lineage | lineage |
| MET | methylation | methylation |
| MISL | mislocation | mislocation |
| MUT | mutation | activating mutation missense |
| NC-EXP | nuclear expression | nuclear expression |
| OEXP | overexpression | overexpression |
| ONCO-MUT | oncogenic mutation | oncogenic mutation |
| PHOS | phosphorylation | phosphorylation |
| PLM | polymorphism | polymorphism |
| PWD | pathway down-regulation | PWD |
| PWU | pathway up-regulation | PWU |
| SKP-MUT | skipping mutation | skipping mutation |
| SLN | sublineage | sublineage |
| SNV | simple nucleotide variation | snv |
| SPAV | splice acceptor variant | splice acceptor variant |
| SPDV | splice donor variant | splice donor variant |
| SPV | splice variant | splice variant; splice site insertion |
| TC-MUT | truncation mutation | truncation mutation |
| UDEF | gene | gene |
| UEXP | underexpression | underexpression |
| WT | wild type | wild type |

**Table S2.**

| **Normalized** **Attribute** | **Specification** |
| --- | --- |
| Tissue type | The resources that iGMDR integrates included different databases and literatures, and their names for cancer types are diverse. To this end, we introduced oncotree (CMO Tumor Type Tree, http://www.oncokb.org/) to manually calibrate all cancer types, promoting structured data storage and use. |
| Cancer type | Oncotree classifies cancer types and associates them with cancer tissue. To facilitate model-related information based on the application of cancer tissue, we involved tissue information in iGMDR. It's worth mentioning that none of the original data we collected provided information on the type of cancer tissue. |
| Drug name | In order to manage data generally, we manually standardize the drug descriptions with DrugBank (Law, et al., 2014) and PubChem (Kaiser, 2005). |
| Gene | All gene names refer to HGNC (https://www.genenames.org/). In addition, we used gene ID (NCBI Entrez ID) as the index to facilitate data processing in the database table. |
| Model description | Logic “and” (denoted as &), which represents the union of two single characteristics; logic “or” (denoted as \|), which indicates that two characteristics can be replaced with each other; and logical “not” (denoted as ¬), which indicates that relevant characteristics are not detected in tumor cell. |

**Table S3.**

| **Interactive Attribute** | **Correlation** | **External Resource** |
| --- | --- | --- |
| Composition | Drug | PubChem, ChEMBL, LINCS (Koleti, et al., 2018) |
| Structure |  | PubChem |
| Target |  | TTD (Li, et al., 2018), DrugBank, GDSC, CTRP, CCLE |
| Signalling Pathway |  | KEGG, CTRP, GDSC |
| Class |  | DrugBank |
| Summary | Gene | MyGene API (Xin, et al., 2016), OMIM, PHARMGKB (Whirl-Carrillo, et al., 2012) |
| Function |  | GO |
| Signalling Pathway |  | KEGG, PHARMGKB, Wikipathways, Smpdb, Reactome |
| Expression |  | NCBI BioProject |

**Table S4.**

| **Pathway ID** | **Description** | **Model Num.** | **Drug Num.** |
| --- | --- | --- | --- |
| WP4172 | PI3K-Akt signaling Pathway | 12 | 3 |
| hsa04150 | mTOR signaling pathway | 11 | 3 |
| hsa04152 | AMPK signaling pathway | 7 | 2 |
| R-HSA-5633007 | Regulation of TP53 activity | 10 | 4 |
